# Beat gestures influence which speech sounds you hear

**DOI:** 10.1101/2020.07.13.200543

**Authors:** Hans Rutger Bosker, David Peeters

**Affiliations:** Max Planck Institute for Psycholinguistics, PO Box 310, 6500 AH, Nijmegen, The Netherlands; Donders Institute for Brain, Cognition and Behaviour, Radboud University, Nijmegen, the Netherlands; Tilburg University, Department of Communication and Cognition, TiCC, Tilburg, the Netherlands

**Keywords:** beat gestures, lexical stress, vowel length, speech perception, manual McGurk effect

## Abstract

Beat gestures – spontaneously produced biphasic movements of the hand – are among the most frequently encountered co-speech gestures in human communication. They are closely temporally aligned to the prosodic characteristics of the speech signal, typically occurring on lexically stressed syllables. Despite their prevalence across speakers of the world’s languages, how beat gestures impact spoken word recognition is unclear. Can these simple ‘flicks of the hand’ influence speech perception? Across six experiments, we demonstrate that beat gestures influence the explicit and implicit perception of lexical stress (e.g., distinguishing *OBject* from *obJECT*), and in turn, can influence what vowels listeners hear. Thus, we provide converging evidence for a *manual McGurk effect*: even the simplest ‘flicks of the hands’ influence which speech sounds we hear.

**SIGNIFICANCE STATEMENT:** Beat gestures are very common in human face-to-face communication. Yet we know little about their behavioral consequences for spoken language comprehension. We demonstrate that beat gestures influence the explicit and implicit perception of lexical stress, and, in turn, can even shape what vowels we think we hear. This demonstration of a *manual McGurk effect* provides some of the first empirical support for a recent multimodal, situated psycholinguistic framework of human communication, while challenging current models of spoken word recognition that do not yet incorporate multimodal prosody. Moreover, it has the potential to enrich human-computer interaction and improve multimodal speech recognition systems.

## INTRODUCTION

The human capacity to communicate evolved primarily to allow for efficient face-to-face interaction ^1^. As humans, we are indeed capable of translating our thoughts into communicative messages, which in everyday situations often comprise concurrent auditory (speech) and visual (lip movements, facial expressions, hand gestures, body posture) information ^2–4^. Speakers thus make use of different channels (mouth, eyes, hands, torso) to get their messages across, while addressees have to quickly and efficiently integrate or *bind* the different ephemeral signals they concomitantly perceive ^2,5,6^. This process is skilfully guided by top-down predictions ^7–9^ that exploit a life-time of communicative experience, acquired world knowledge, and personal common ground built up with the speaker ^10–12^.

The classic McGurk effect ^13^ is an influential – though not entirely uncontroversial ^14–16^ – illustration of how visual input influences the perception of speech sounds. In a seminal study ^13^, participants were made to repeatedly listen to the sound /ba/. At the same time, they watched the face of a speaker in a video whose mouth movements corresponded to the sound /ga/. When asked what they heard, a majority of participants reported actually perceiving the sound /da/. This robust finding indicates that, during language comprehension, our brains take both verbal (here: auditory) and non-verbal (here: visual) aspects of communication into account ^17,18^, providing the addressee with a best guess of what exact message a speaker intends to convey. We thus listen not only with our ears, but also with our eyes.

Visual aspects of everyday human communication are not restricted to subtle mouth or lip movements. In close coordination with speech, the *hand gestures* we spontaneously and idiosyncratically make during conversations help us express our thoughts, emotions, and intentions^19–21^. We manually point at things to guide the attention of our addressee to relevant aspects of our immediate environment ^22–25^, enrich our speech with iconic gestures that in handshape and movement resemble what we are talking about ^26–28^, and produce simple flicks of the hand aligned to the acoustic prominence of our spoken utterance to highlight relevant points in speech ^29–34^. Over the past decades, it has become clear that such co-speech manual gestures are semantically aligned and temporally synchronized with the speech we produce ^35–40^. It is an open question, however, whether the temporal synchrony between these hand gestures and the speech signal can actually influence which speech sounds we hear.

In this study, we test for the presence of such a potential *manual* McGurk effect. Do hand gestures, like observed lip movements, influence what exact speech sounds we perceive? We here focus on manual *beat gestures* – commonly defined as biphasic (e.g., up and down) movements of the hand that speakers spontaneously produce to highlight prominent aspects of the concurrent speech signal ^21,31,41^. Beat gestures are amongst the most frequently encountered gestures in naturally occurring human communication ^21^ as the presence of an accompanying beat gesture allows for the enhancement of the perceived prominence of a word ^31,42,43^. Not surprisingly, they hence also feature prominently in politicians’ public speeches ^44,45^. By highlighting concurrent parts of speech and enhancing its processing ^46–49^, beat gestures may even increase memory recall of words in both adults ^50^ and children ^51–53^. As such, they form a fundamental part of our communicative repertoire and also have direct practical relevance.

In contrast to iconic gestures, beat gestures are not related to the spoken message on a semantic level, but only on a temporal level: their apex is aligned to vowel onset in lexically stressed syllables carrying prosodic emphasis ^20,21,34,54^. Neurobiological studies suggest that listeners may tune oscillatory activity in the alpha and theta bands upon observing the initiation of a beat gesture to anticipate processing an assumedly important upcoming word ^44^. Bilateral temporal areas of the brain may in parallel be activated for efficient integration of beat gestures and concurrent speech^48^. However, evidence for behavioral effects of temporal gestural alignment on speech perception remains scarce. Given the close temporal alignment between beat gestures and lexically stressed syllables in speech production, ideal observers ^55^ could in principle use beat gestures as a cue to *lexical stress* in perception.

Lexical stress is indeed a critical speech cue in spoken word recognition. It is well established that listeners commonly use suprasegmental spoken cues to lexical stress, such as greater amplitude, higher fundamental frequency (F0), and longer syllable duration, to constrain online lexical access, speeding up word recognition, and increasing perceptual accuracy ^56,57^. Listeners also use visual articulatory cues on the face to perceive lexical stress when the auditory signal is muted ^58–60^. Nevertheless, whether non-facial visual cues, such as beat gestures, also aid in lexical stress perception, and might even do so in the presence of auditory cues to lexical stress – as in everyday conversations – is unknown. These questions are relevant for our understanding of face-to-face interaction, where multimodal cues to lexical stress might considerably increase the robustness of spoken communication, for instance in noisy listening conditions ^61^, and even when the auditory signal is clear ^62^.

Indeed, a recent theoretical account proposed that *multimodal low-level integration* must be a general cognitive principle fundamental to our human communicative capacities ^2^. However, evidence showing that the integration of information from two separate communicative modalities (speech, gesture) modulates the mere perception of information derived from one of these two modalities (e.g., speech) is lacking. For instance, earlier studies have failed to find effects of simulated pointing gestures on lexical stress perception, leading to the idea that manual gestures provide information only about sentential emphasis; not about relative emphasis placed on individual syllables within words ^63^. As a result, there is no formal account or model of how audiovisual prosody might help spoken word recognition ^64–66^. Hence, demonstrating a manual McGurk effect would imply that existing models of speech perception need to be fundamentally expanded to account for how multimodal cues influence auditory word recognition.

Below, we report the results of four experiments (and two direct replications: Experiments S1-S2) in which we tested for the existence and robustness of a manual McGurk effect. The experiments each approach the theoretical issue at hand from a different methodological angle, varying the degree of explicitness of the participants’ experimental task and their response modality (perception *vs.* production). Experiment 1 uses an explicit task to test whether observing a beat gesture influences the perception of *lexical stress* in disyllabic spoken stimuli. Using the same stimuli, Experiment 2 *implicitly* investigates whether seeing a concurrent beat gesture influences how participants vocally shadow (i.e., repeat back) an audiovisual speaker’s production of disyllabic stimuli. Experiment 3 then uses those shadowed productions as stimuli in a perception experiment to implicitly test whether naïve listeners are biased towards perceiving lexical stress on the first syllable if the shadowed production was produced in response to an audiovisual stimulus with a beat gesture on the first syllable. Finally, Experiment 4 tests whether beat gestures may influence what *vowels* (long *vs*. short) listeners perceive. The perceived prominence of a syllable influences the expected duration of that syllable’s vowel nucleus ^i.e., longer if stressed, 67^. Thus, if listeners use beat gestures as a cue to lexical stress, they may perceive a vowel midway between Dutch short /ɑ/ and long /a:/ as /ɑ/ when presented with a concurrent beat gesture, but as /a:/ without it.

For all experiments, we hypothesized that, if the manual McGurk effect exists, we should see effects of the presence and timing of beat gestures on the perception and production of lexical stress (Experiments 1-3) and, in turn, on vowel perception (Experiment 4). In short, we found evidence in favor of a manual McGurk effect in all four experiments and in both direct replications: the perception of lexical stress and vowel identity is consistently influenced by the temporal alignment of beat gestures to the speech signal.

## RESULTS

Experiment 1 tested whether beat gestures influence how listeners explicitly categorize disyllabic spoken stimuli as having initial stress or final stress (e.g., *WAsol* vs. *waSOL*; capitals indicate lexical stress). Native speakers of Dutch were presented with an audiovisual speaker (facial features masked in the actual experiment to avoid biasing effects of articulatory cues) producing non-existing pseudowords in a sentence context: *Nu zeg ik het woord… [target]* “Now say I the word… [target]”. Pseudowords were manipulated along a 7-step lexical stress continuum, varying F0 in opposite directions for the two syllables, thus ranging from a ‘strong-weak’ (step 1) to a ‘weak-strong’ prosodic pattern (step 7). Critically, the audiovisual speaker always produced a beat gesture whose apex was manipulated to be either aligned to the onset of the first vowel or the second vowel by varying the silent interval preceding the pseudoword (cf. Figure 1A). The task involved two-alternative forced choice (2AFC) between the pseudoword having stress on the first or second syllable. For comparison, in another block, the same participants also categorized audio-only versions of the stimuli (block order counter-balanced). Outcomes of a Generalized Linear Mixed Model with a logistic linking function ^GLMM; 68^ demonstrated that participants, in the audiovisual block, were significantly more likely to report hearing stress on the first syllable if the beat gesture was aligned to the onset of the first vowel (main effect of Beat Condition: *β* = 1.142 in logit space, *p* < 0.001), as shown by the difference between the blue and orange solid lines in Figure 1B (mean difference in proportions between the two Beat Conditions: 0.20). No such effect was observed in the audio-only condition (interaction Beat Condition x Modality: *β* = −0.813, *p* < 0.001). These findings demonstrate that the temporal alignment of beat gestures to the speech signal influences the explicit perception of lexical stress.

**Figure 1.**
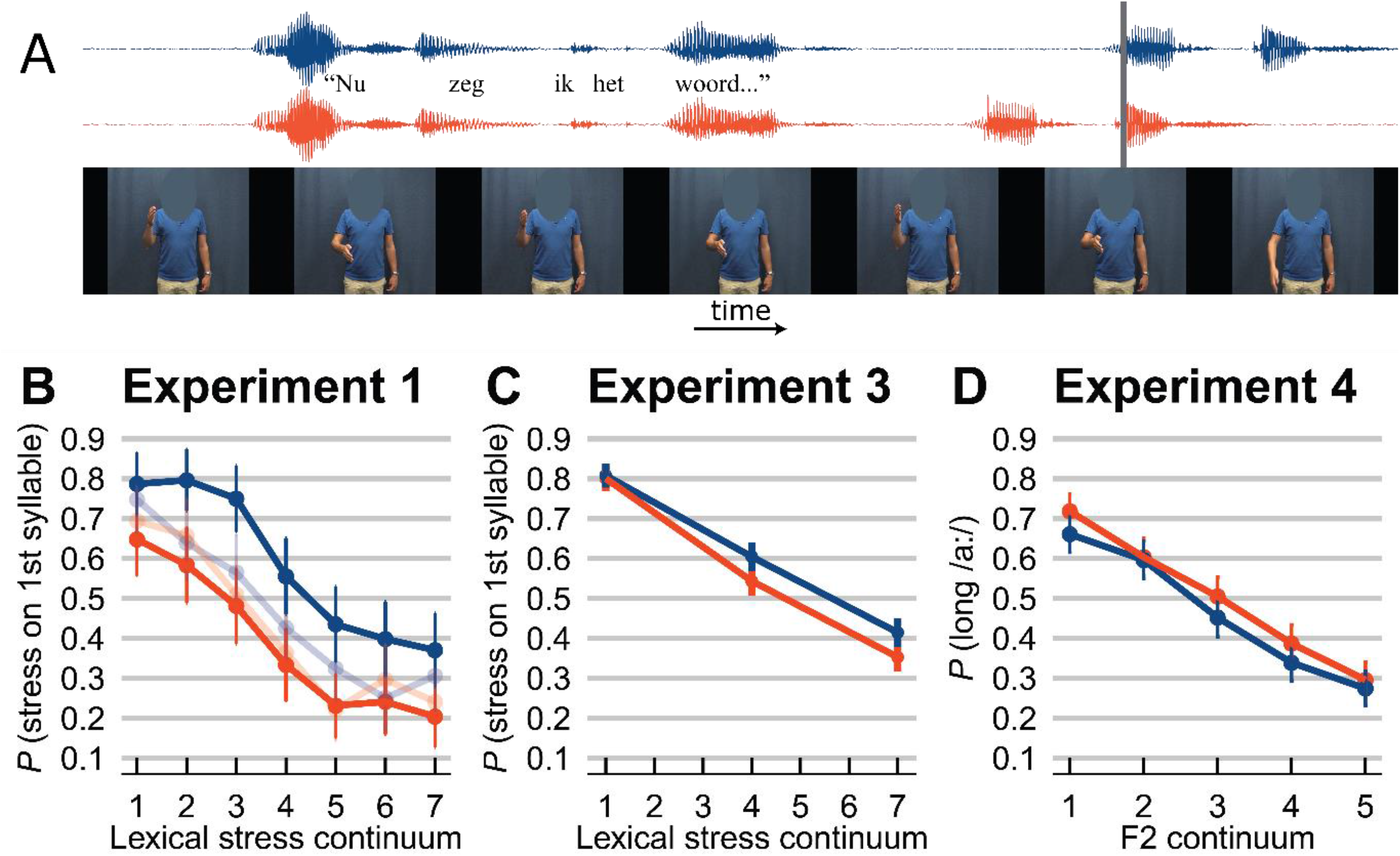
Example audiovisual stimulus and results from Experiments 1, 3, and 4. A. In Experiments 1, 2, and 4, participants were presented with an audiovisual speaker (facial features masked in the actual experiment) producing the Dutch carrier sentence *Nu zeg ik het woord…* “Now say I the word…” followed by a disyllabic target pseudoword (e.g., *bagpif*). The speaker produced three beat gestures, with the last beat gesture’s apex falling in the target time window. Two auditory conditions were created by aligning the onset of either the first (top waveform in blue) or the second vowel of the target pseudoword (bottom waveform in orange) to the final gesture’s apex (gray vertical line). **B.** Participants in Experiment 1 categorized audiovisual pseudowords as having lexical stress on either the first or the second syllable. Each target pseudoword was sampled from a 7-step continuum (on x-axis) varying F0 in opposite directions for the two syllables. In the audiovisual block (solid lines), participants were significantly more likely to report perceiving lexical stress on the first syllable if the speaker produced a beat gesture on the first syllable (in blue) vs. second syllable (in orange). No such difference was observed in the audio-only control block (transparent lines). **C.** Participants in Experiment 3 categorized the audio-only shadowed productions, recorded in Experiment 2, as having lexical stress on either the first or the second syllable. If the audiovisual stimulus in Experiment 2 involved a pseudoword with a beat gesture on the first syllable, the elicited shadowed production was more likely to be perceived as having lexical stress on the first syllable (in blue). Conversely, if the audiovisual stimulus in Experiment 2 involved a pseudoword with a beat gesture on the second syllable, the elicited shadowed production was more likely to be perceived as having lexical stress on the second syllable (in orange). **D.** Participants in Experiment 4 categorized audiovisual pseudowords as containing either short /ɑ/ or long /a:/ (e.g., *bagpif* vs. *baagpif*). Each target pseudoword contained a first vowel that was sampled from a 5-step F2 continuum from long /a:/ to short / ɑ/ (on x-axis). Moreover, prosodic cues to lexical stress (F0, amplitude, syllable duration) were set to ambiguous values. If the audiovisual speaker produced a beat gesture on the first syllable (in blue), participants were biased to perceive the first syllable as stressed, making the ambiguous first vowel relatively short for a stressed syllable, leading to a lower proportion of long /a:/ responses. Conversely, if the audiovisual speaker produced a beat gesture on the second syllable (in orange), participants were biased to perceive the initial syllable as unstressed, making the ambiguous first vowel relatively long for an unstressed syllable, leading to a higher proportion of long /a:/ responses. For all panels, error bars enclose 1.96 x SE on either side; that is, the 95% confidence intervals.

However, the task in Experiment 1, asking participants to categorize the audiovisual stimuli as having stress on either the first or second syllable, is very explicit and presumably susceptible to response strategies. Therefore, Experiment 2 tested lexical stress perception *implicitly* and in another response modality by asking participants to overtly shadow the audiovisual pseudowords from Experiment 1. To reduce the load on the participants, we selected only three relevant steps from the lexical stress continua: two relatively clear steps (1 and 7) and one relatively ambiguous step (step 4). Critically, lexical stress was never mentioned in any part of the instructions. We hypothesized that, if participants saw the audiovisual speaker produce a beat gesture on the second syllable, the suprasegmental characteristics of their shadowed productions would resemble a ‘weak-strong’ prosodic pattern (i.e., greater amplitude, higher F0, longer duration on the second syllable). Acoustic analysis of the amplitude, F0, and duration measurements of the shadowed productions by three separate Linear Mixed Models (one for each acoustic cue) showed small but significant effects of Beat Condition, visualized in Figure 2. That is, if the beat gesture was aligned to the second syllable, participants presumably implicitly perceived lexical stress on the second syllable, and therefore their own shadowed second syllables were produced with greater amplitude and a longer duration, while their first syllables were shorter.

**Figure 2.**
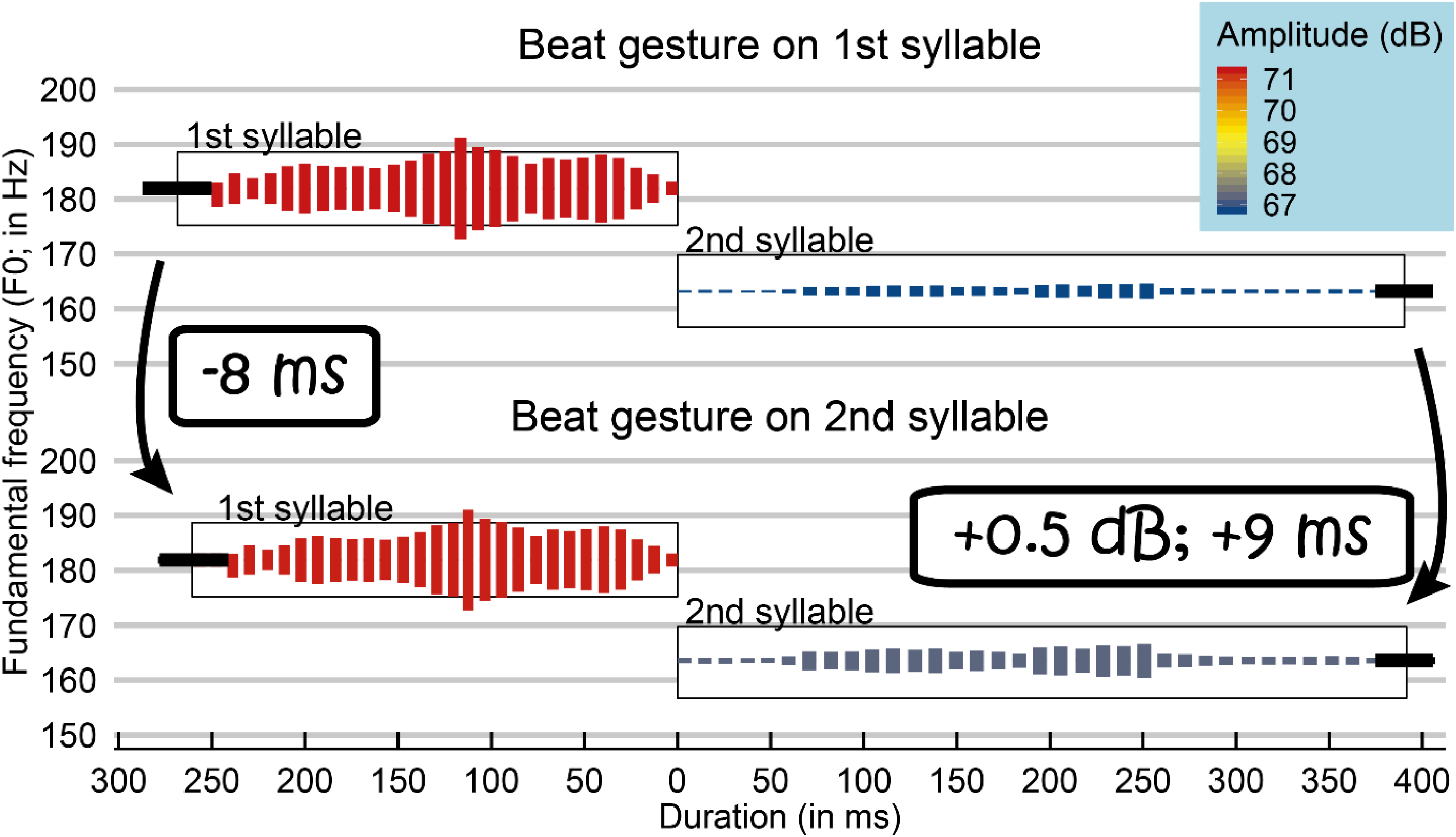
Predicted estimates from the combined models of Experiment 2. In Experiment 2, participants shadowed the audiovisual sentence-final pseudowords in either Beat Condition. The figure shows stylized wave forms for the two syllables of the shadowed productions, on either side of the zero point on the x-axis, showing their durations (ms), aligned around the onset of the second syllable (at 0). The bold horizontal error bars at the extremes of the stylized waveforms indicate duration SE. The vertical position of the (black rectangles surrounding the) stylized waveforms shows the produced F0 (in Hz on y-axis; height of rectangles indicates F0 SE on either side of the midpoint). The color gradient and height of the stylized waveforms indicates average amplitude (dB). The overall differences between first and second syllables show pronounced word-final effects: the last syllable of a word typically has a longer duration, and a lower F0 and amplitude. Critically, after audiovisual trials with a beat gesture on the second syllable (lower panel; compared to a beat gesture on the first syllable in the upper panel), shadowers produced their first syllables with a shorter duration and their second syllables with a longer duration and higher amplitude. No effect of F0 was observed.

Note that the effects of beat gestures in Experiment 2 were robust but small. Furthermore, no effect of Beat Condition was found on the F0 of the shadowed productions, the primary cue to stress in Dutch ^69^. Therefore, Experiment 3 tested the *perceptual relevance* of the acoustic adjustments participants made to their shadowed productions in response to the two beat gesture conditions. Each participant listened to all shadowed productions from two shadowers from Experiment 2 and indicated whether they perceived the shadowed productions to have stress on the first or second syllable (2AFC). Critically, this involved audio-only stimuli and the participants in Experiment 3 were unaware of the beat gesture manipulation in Experiment 2. Nevertheless, outcomes of a GLMM on the binomial categorization data demonstrated that participants were significantly biased to report hearing stress on the first syllable of the audio-only shadowed productions if the audiovisual stimulus that elicited those shadowed productions had a beat gesture on the first syllable (main effect of Beat Condition: *β* = 0.232 in logit space, *p* = 0.003). This is indicated by the difference between the blue and orange lines in Figure 1C (mean difference in proportions between the two Beat Conditions: 0.05). Moreover, an identical replication of this experiment, Experiment S1 (reported in the Supplementary Material), reported similar findings using the same stimuli in a new participant sample. These findings suggest that the acoustic adjustments in response to the different beat gesture alignments in Experiment 2 were perceptually salient enough for new participants to notice them and use them as cues to lexical stress identification. Thus, it provides supportive evidence that beat gestures influence *implicit* lexical stress perception.

Finally, Experiment 4 tested the strongest form of a manual McGurk effect: can beat gestures influence the perceived identity of *individual speech sounds*? The rationale behind Experiment 4 is grounded in the observation that listeners take a syllable’s perceived prominence into account when estimating its vowel’s duration ^67,70^. Vowels in stressed syllables typically have a longer duration than vowels in unstressed syllables. Hence, in perception, listeners expect a vowel in a stressed syllable to have a longer duration. That is, one and the same vowel duration can be perceived as relatively short in one (stressed) syllable, but as relatively long in another (unstressed syllable). In languages where vowel duration is a critical cue to phonemic contrasts (e.g., to coda stop voicing in English, e.g., “coat - code” ^70^; to vowel length contrasts in Dutch, e.g., *tak* /tɑk/ “branch” - *taak* /ta:k/ “task” ^71^), these expectations can influence the perceived identity of speech sounds.

Participants in Experiment 4 were presented with audiovisual stimuli, much like those used in Experiments 1-2. However, we used a new set of disyllabic pseudowords whose initial vowel was manipulated to be ambiguous between Dutch /ɑ-a:/ in its duration while varying the second formant frequency (F2) in 5-step F2 continua, which is a secondary cue to the /ɑ-a:/ contrast in Dutch ^71^ but not a cue to lexical stress. Moreover, each pseudoword token was manipulated to be ambiguous with respect to its suprasegmental cues to lexical stress (set to average values of F0, amplitude, and duration). Taken together, these manipulations resulted in pseudowords that were ambiguous in lexical stress and vowel duration, but varied in F2 as cue to vowel identity. Participants indicated whether they heard an /ɑ/ or /a:/ as initial vowel of the pseudoword (2AFC; e.g., *bagpif* or *baagpif*?); lexical stress was never mentioned in any part of the instructions. Outcomes of a GLMM on the binomial vowel categorization data demonstrated an effect of Beat Condition on participants’ vowel perception (*β* = −0.240 in logit space, *p* = 0.023). That is, if the beat gesture was aligned to the first syllable (blue line in Figure 1D), participants were more likely to perceive the first syllable as stressed, rendering the ambiguous vowel duration in the first syllable as relatively short for a stressed syllable, lowering the proportion of long /a:/ responses.

Conversely, if the beat gesture was aligned to the second syllable (orange line in Figure 1D), participants were more likely to perceive the first syllable as unstressed, making the ambiguous vowel duration in the first syllable relatively long for an unstressed syllable, leading to a small increase in the proportion of long /a:/ responses (mean difference in proportions between the two Beat Conditions: 0.04). This finding, corroborated by similar observations in an identical replication of this experiment, Experiment S2 (see Supplementary Material), showcases the strongest form of the manual McGurk effect: beat gestures influence which speech sounds we hear.

## DISCUSSION

To what extent do we listen with our eyes? The classic McGurk effect illustrates that listeners take both visual (lip movements) and auditory (speech) information into account to arrive at a best possible approximation of the exact message a speaker intends to convey ^13^. Here we provide evidence in favor of a *manual* McGurk effect: the beat gestures we see influence what exact speech sounds we hear. Specifically, we observed in four experiments that beat gestures influence both the explicit and implicit perception of lexical stress, and in turn may influence the perception of individual speech sounds. To our understanding, this is the first demonstration of behavioral consequences of beat gestures on low-level speech perception. These findings support the idea that everyday language comprehension is primordially a multimodal undertaking in which the multiple streams of information that are concomitantly perceived through the different senses are quickly and efficiently integrated by the listener to make a well-informed best guess of what specific message a speaker is intending to communicate.

The four experiments we presented made use of a combination of implicit and explicit paradigms. Not surprisingly, the *explicit* nature of the task used in Experiment 1 led to strong effects of the timing of a perceived beat gesture on the lexical stress pattern participants reported to hear. Although the *implicit* effects in subsequent Experiments 2-4 were smaller, conceivably partially due to suboptimal control over the experimental settings in online testing, they were nevertheless robust and replicable (cf. Experiments S1-S2 in Supplementary Material). Moreover, these beat gesture effects even surfaced when the auditory cues were relatively clear (i.e., at the extreme ends of the acoustic continua), suggesting that these effects also prevail in naturalistic face-to-face communication outside the lab. In fact, the naturalistic nature of the audiovisual stimuli would seem key to the effect, since the temporal alignment of artificial hand movements (i.e., an animated line drawing of a disembodied hand) does not influence lexical stress perception ^see 63^. Perhaps the present study even underestimates the behavioral influence of beat gestures on speech perception in everyday conversations, since all auditory stimuli in the experiments involved ‘clean’ (i.e., noise-free) speech. However, in real life situations, the speech of an attended talker oftentimes occurs in the presence of other auditory signals, such as background music, competing speech, or environmental noise; in such listening conditions, the contribution of visual beat gestures to successful communication would conceivably be even greater.

The outcomes of the present study challenge current models of (multimodal) word recognition, which would need to be extended to account for how prosody, specifically multimodal prosody, supports and modulates word recognition. Audiovisual prosody is implemented in the fuzzy logical model of perception ^72^, but how this information affects lexical processing is underspecified. Moreover, Experiment 4 implicates that the multimodal suprasegmental and segmental ‘analyzers’ would have to allow for interaction, since one can influence the other. Therefore, existing models of audiovisual speech perception and word recognition thus need to be expanded to account for (1) how suprasegmental prosodic information is processed ^73^; (2) how suprasegmental information interacts with and modulates segmental information ^74^; and (3) how multimodal cues influence auditory word recognition ^17,58^.

A recent, broader theoretical account of human communication (i.e., beyond speech alone) proposed that two domain-general mechanisms form the core of our ability to understand language: multimodal low-level integration and top-down prediction ^2^. The present results are indeed in line with the idea that *multimodal low-level integration* is a general cognitive principle supporting language understanding. More precisely, we here show that this mechanism operates across communicative modalities. The cognitive and perceptual apparatus that we have at our disposal apparently not just binds auditory and visual information that derive from the same communicative channel (i.e., articulation), but also combines visual information derived from the hands with concurrent auditory information – crucially to such an extent that what we hear is modulated by the timing of the beat gestures we see. As such, what we perceive is the model of reality that our brains provide us by binding visual and auditory communicative input, and not reality itself.

In addition to multimodal low-level integration, earlier research has indicated that also *top-down predictions* play a pivotal role in the perception of co-speech beat gestures. It seems reasonable to assume that listeners throughout life build up the implicit knowledge that beat gestures are commonly tightly paired with acoustically prominent verbal information. As the initiation of a beat gesture typically slightly precedes the prominent acoustic speech feature ^40^, these statistical regularities are then indeed used in a top-down manner to allocate additional attentional resources to an upcoming word ^44^.

The current first demonstration of the influence of simple ‘flicks of the hands’ on low-level speech perception should be considered an initial step in our understanding of how bodily gestures in general may affect speech perception. Future research may investigate whether the temporal alignment of types of non-manual gestures, such as head nods and eyebrow movements, also impact what speech sounds we hear. Moreover, other behavioral consequences beyond lexical stress perception could be implicated, ranging from phase-resetting neural oscillations ^44^ with consequences for speech intelligibility and phoneme identification ^75,76^, to facilitating selective attention in ‘cocktail party’ listening ^77–79^, word segmentation ^80^, and word learning ^81^.

Finally, the current findings have several practical implications. Speech recognition systems may be improved when including multimodal signals, even beyond articulatory cues on the face. Presumably beat gestures are particularly valuable multimodal cues, because, while cross-language differences exist in the use and weighting of acoustic suprasegmental prosody ^e.g., amplitude, F0, duration, 82–84^, beat gestures could presumably function as a language-universal prosodic cue. Also, the field of human-computer interaction could benefit from the finding that beat gestures support spoken communication, for instance by enriching virtual agents and avatars with human-like gesture-to-speech alignment. The use of such agents in immersive virtual reality environments will in turn allow for taking the study of language comprehension into naturalistic, visually rich and dynamic settings while retaining the required degree of experimental control ^85^. By acknowledging that spoken language comprehension involves the integration of auditory sounds, visual articulatory cues, and even the simplest of manual gestures, it will be possible to significantly further our understanding of the human capacity to communicate.

## MATERIALS AND METHODS

### Participants

Native speakers of Dutch were recruited from the Max Planck Institute’s participant pool (Experiment 1: *N* = 20; 14 females (f), 6 males (m); *M*_age_ = 25, range = 20-34; Experiment 2: *N* = 26; 22f, 4m; *M*_age_ = 23, range = 19-30; Experiment 3: *N* = 26; 25f, 1m; *M*_age_ = 24, range = 20-28; Experiment 4 was performed by the same individuals as those in Experiment 3). All gave informed consent as approved by the Ethics Committee of the Social Sciences department of Radboud University (project code: ECSW2014-1003-196). All research was performed in accordance with relevant guidelines and regulations. Participants had normal hearing, had no speech or language disorders, and took part in only one of our experiments.

## Materials

### Experiments 1 & 2

For our auditory target stimuli, we used 12 disyllabic pseudowords that are phonotactically legal in Dutch (e.g., *wasol* /ʋa.sɔl/; see Table S1 in Supplementary Material). In Dutch, lexical stress is cued by three suprasegmental cues: amplitude, fundamental frequency (F0), and duration ^69^. Since, in Dutch, F0 is a primary cue to lexical stress in perception, we used 7-step lexical stress continua – adopted from an earlier student project ^86^ – varying in mean F0 while keeping amplitude and duration cues constant, thus ranging from a ‘strong-weak’ prosodic pattern (SW; step 1) to a ‘weak-strong’ prosodic pattern (WS; step 7).

These continua had been created by recording an SW (i.e., stress on initial syllable) and a WS version (i.e., stress on last syllable) of each pseudoword from the first author, a male native speaker of Dutch. We first measured the average duration and amplitude values, separately for the first and second syllable, across the stressed and unstressed versions: mean duration syllable 1 = 202 ms; syllable 2 = 395 ms; mean amplitude syllable 1 = 68.1 dB; syllable 2 = 64 dB. Then, using Praat ^87^, we set the duration and amplitude values of the two syllables of each pseudoword to these ambiguous values. Subsequently, F0 was manipulated along a 7-step continuum, with the two extremes and the step size informed by the talker’s originally produced F0. Manipulations were always performed in an inverse manner for the two syllables of each pseudoword: while the mean F0 of the first syllable decreased along the continuum (from 159.2 to 110.6 Hz in steps of 8.1 Hz), the mean F0 of the second syllable increased (from 95.7 to 141.3 Hz in steps of 7.6 Hz). Moreover, rather than setting the F0 within each syllable to a fixed value, more natural output was created by including a fixed F0 declination within the first syllable (linear decrease of 23.5 Hz) and second syllable (38.2 Hz). Pilot data from a categorization task on these manipulated stimuli showed that these manipulations resulted in 7-step F0 continua that appropriately sampled the SW-to-WS perceptual space (i.e., from 0.91 proportion SW responses for step 1 to 0.16 proportion SW responses for step 7).

To create audiovisual stimuli, the first author was video-recorded using a Canon XF205 camera (50 frames per second; resolution: 1280 by 720 pixels) with an external Sennheiser ME64 directional microphone (audio sampling frequency: 48 kHz). Recordings were made from knees up on a neutral background in the Gesture Lab at the Max Planck Institute for Psycholinguistics. The speaker produced the Dutch carrier sentence *Nu zeg ik het woord… kanon* “Now say I the word… cannon”, with lexical stress on the second syllable of *kanon*. Concurrently, he produced three beat gestures in this sentence, with their apex aligned to the syllables *Nu*, *woord*, and *–non*. The manipulated auditory pseudowords described above were combined with this audiovisual recording using *ffmpeg* (version 4.0; available from http://ffmpeg.org/) by (1) removing the original *kanon* target word, (2) inserting a silent interval, and (3) inserting the manipulated pseudowords (cf. Figure 1A). By modulating the length of the intervening silent interval, the onset of either the first or the second vowel of the target pseudoword was aligned to the final beat gesture’s apex. Furthermore, the facial features of the talker were masked to hide any visual articulatory cues to lexical stress. Finally, in order to mask the cross-spliced nature of the audio, combining recordings with variable room acoustics, the silent interval and pseudoword were mixed with stationary noise from a silent recording of the Gesture Lab.

### Experiment 4

We created 8 new disyllabic pseudowords, with either a short /ɑ/ or a long /a:/ as first vowel (e.g., *bagpif* /bɑx.pɪf/ vs. *baagpif* /ba:x.pɪf/; cf. Table S1). This new set had a fixed syllable structure (CVC.CVC, only stops and fricatives) to reduce item variance and to facilitate syllable-level annotations. The same male talker as before was recorded producing these pseudowords in four versions: with short /ɑ/ and stress on the first syllable (e.g., *BAGpif*); with short /ɑ/ and stress on the second syllable (*bagPIF*); with long /a:/ and stress on the first syllable (*BAAGpif*); with long /a:/ and stress on the second syllable (*baagPIF*).

After manual annotation of syllable onsets and offsets, we measured the values of the suprasegmental cues to lexical stress (F0, amplitude, and duration) in all four conditions. Pseudowords with long /a:/ and stress on the first syllable were selected for manipulation. First, we manipulated the cues to lexical stress by setting these to ambiguous values using PSOLA in Praat: each first syllable was given the same mean value calculated across all stressed and unstressed first syllables (mean F0 = 154 Hz; original contour maintained; amplitude = 65.88 dB; duration = 263 ms), and each second syllable was given the same mean value across all stressed and unstressed second syllables (mean F0 = 159 Hz; original contour maintained; amplitude = 63.50 dB; duration = 414 ms). This resulted in manipulated pseudowords that were ambiguous in their prosodic cues to lexical stress. Since this included manipulating duration across all recorded conditions as well, it also meant that the duration cues to the identity of the first vowel were ambiguous.

Second, the first /a:/ vowel was extracted and manipulated to form a spectral continuum from long /a:/ to short /ɑ/. In Dutch, the /ɑ-a:/ vowel contrast is cued by both spectral (lower first (F1) and second formant (F2) values for /ɑ/) and temporal cues ^shorter duration for /ɑ/, 71^. We decided to create F2 continua since F2 is no cue to lexical stress in Dutch (in fact, in our original recordings F2 values did not differ between stressed and unstressed conditions) while varying F2 does influence vowel quality perception. These spectral manipulations were based on Burg’s LPC method in Praat, with the source and filter models estimated automatically from the selected vowel. The formant values in the filter models were adjusted to result in a constant F1 value (750 Hz, ambiguous between /ɑ/ and /a:/; value based on the original recordings) and 5 F2 values (step 1 = 1325 Hz, step 5 = 1125 Hz, step size = 50 Hz; values based on the original recordings). Then, the source and filter models were recombined. Finally, the manipulated vowel tokens were spliced into the pseudowords. Taken together, these manipulations resulted in pseudoword tokens that were ambiguous in lexical stress (average values of F0, amplitude, and duration), but varied in F2 as cue to vowel identity.

To create audiovisual stimuli, these manipulated pseudowords were spliced into the audiovisual stimuli from Experiment 1. Once again, manipulating the silent interval between carrier sentence offset and target onset resulted in two different alignments of the last beat gesture to either first vowel onset or second vowel onset. Like in Experiment 1, facial features of the talker were masked, and silent intervals as well as manipulated pseudowords were mixed with stationary noise to match the room acoustics across the sentence stimuli.

### Procedure

#### Experiment 1

Participants were tested individually in a sound-conditioning booth. They were seated at a distance of approximately 60 cm in front of a 50,8 cm by 28,6 cm screen and listened to stimuli at a comfortable volume through headphones. Stimulus presentation was controlled by Presentation software (v16.5; Neurobehavioral Systems, Albany, CA, USA).

Participants were presented with two blocks of trials: an audiovisual block, in which the video and the audio content of the manipulated stimuli were concomitantly presented, and an audio-only control block, in which only the audio content but no video content was presented. Block order was counter-balanced across participants. On each trial, the participants’ task was to indicate whether the sentence-final pseudoword was stressed on the first or the last syllable (2-alternative forced choice task; 2AFC).

Before the experiment, participants were instructed to always look at the screen during trial presentations. In the audiovisual block, trials started with the presentation of a fixation cross. After 1000 ms, the audiovisual stimulus was presented (video dimensions: 960 x 864 pixels). At stimulus offset, a response screen was presented with two response options on either side of the screen, for instance: *WAsol* vs. *waSOL*; positioning of response options was counter-balanced across participants. Participants were instructed that capitals indicated lexical stress, that they could enter their response by pressing the “Z” key on a regular keyboard for the left option and the “M” key for the right option, and that they had to give their response within 4 s after stimulus offset; otherwise a missing response was recorded. After their response (or at timeout), the screen was replaced by an empty screen for 500 ms, after which the next trial was initiated automatically. The trial structure in the audio-only control block was identical to the audiovisual block, except that instead of the video content a static fixation cross was presented during stimulus presentation.

Participants were presented with 12 pseudowords, sampled from 7-step F0 continua, in 2 beat gesture conditions (onset of first or second vowel aligned to apex of beat gesture), resulting in 168 unique items in a block. Within a block, item order was randomized. Before the experiment, four practice trials were presented to participants to familiarize them with the materials and the task. Participants were given opportunity to take a short break after the first block.

#### Experiment 2

Experiment 2 used the same procedure as Experiment 1, except that only audiovisual stimuli were used. Also, instead of making explicit 2AFC decisions about lexical stress, participants were instructed to simply repeat back the sentence-final pseudoword after stimulus offset, and to hit the “Enter” key to move to the next trial. No mention of lexical stress was made in the instructions. Participants were instructed to always look at the screen during trial presentations.

At the beginning of the experiment, participants were seated behind a table with a computer screen inside a double-walled acoustically isolated booth. Participants wore a circum-aural Sennheiser GAME ZERO headset with attached microphone throughout the experiment to ensure that amplitude measurements were made at a constant level. Participants were presented with 12 pseudowords, sampled from step 1, 4, and 7 of the F0 continua, in 2 beat gesture conditions, resulting in 72 unique items. Each unique item was presented twice over the course of the experiment, leading to 144 trials per participants. Item order was random and item repetitions only occurred after all unique items had been presented. Before the experiment, one practice trial was presented to participants to familiarize them with the stimuli and the task.

#### Experiment 3

Experiment 3 was run online using PsyToolkit ^version 2.6.1;,88^ because of limitations due to the corona virus pandemic. Each participant was explicitly instructed to use headphones and to run the experiment with undivided attention in quiet surroundings. Each participant was presented with all shadowed productions from two talkers from Experiment 2, leading to 13 different lists, with two participants assigned to each list. Stimuli from the same talker were grouped to facilitate talker and speech recognition, and presented in a random order. The task was to indicate after stimulus offset whether the pseudoword had stress on the first or the second syllable (2AFC). Two response options were presented, with capitals indicating stress (e.g., *WAsol* vs. *waSOL*), and participants used the mouse to select their response. No practice items were presented and the experiment was self-timed.

Note that the same participants who performed Experiment 3 also participated in Experiment 4. To avoid explicit awareness about lexical stress influencing their behavior in Experiment 4, these participants always first ran Experiment 4, and only then took part in Experiment 3.

#### Experiment 4

Experiment 4 was run online using Testable (https://testable.org). Each participant was explicitly instructed to use headphones and to run the experiment with undivided attention in quiet surroundings. Before the experiment, participants were instructed to always look and listen carefully to the audiovisual talker and categorize the first vowel of the sentence-final target pseudoword (2AFC). Specifically, at stimulus offset, a response screen was presented with two response options on either side of the screen, for instance: *bagpif* vs. *baagpif*. Participants entered their response by pressing the “Z” key on a regular keyboard for the left option and the “M” key for the right option. The next trial was only presented after a response had been recorded (ITI = 500 ms). No mention of lexical stress was made in the instructions.

Participants were presented with 8 pseudowords, sampled from 5-step F2 continua, in 2 beat gesture conditions (onset of first or second vowel aligned to apex of beat gesture), resulting in 80 unique items in a block. Each participant received two identical blocks with a unique random order within blocks.

### Statistical analysis

#### Experiment 1

Trials with missing data (*n* = 7; 0.1%) were excluded from analysis. Data were statistically analyzed using a Generalized Linear Mixed Model ^GLMM; 68^ with a logistic linking function as implemented in the lme4 library ^version 1.0.5;,89^ in R ^90^. The binomial dependent variable was participants’ categorization of the target as either having lexical stress on the first syllable (SW; e.g., *WAsol*; coded as 1) or the second syllable (WS; e.g., *waSOL*; coded as 0). Fixed effects were Continuum Step (continuous predictor; scaled and centered around the mean), Beat Condition (categorical predictor; deviation coding, with beat gesture on the first syllable coded as +0.5 and beat gesture on the second syllable coded as −0.5), and Block (categorical predictor with the audiovisual block mapped onto the intercept), and all interactions. The use of deviation coding of two-level categorical factors (i.e., coded with −0.5 and +0.5) allows us to test main effects of these predictors, since with this coding the grand mean is mapped onto the intercept. The model also included Participant and Pseudoword as random factors, with by-participant and by-item random slopes for all fixed effects, as advocated by Barr et al. ^91^.

The model showed a significant effect of Continuum Step (*β* = −1.093, *SE* = 0.284, *z* = −3.850, *p* < 0.001), indicating that higher continuum steps led to lower proportions of SW responses. It also showed an effect of Beat Condition (*β* = 1.142, *SE* = 0.186, *z* = 6.155, *p* < 0.001), indicating that – in line with our hypothesis – listeners were biased towards perceiving lexical stress on the first syllable if there was a beat gestures on that first syllable. Since the audiovisual block was mapped onto the intercept, this simple effect of Beat Condition should only be interpreted with respect to the audiovisual block. In fact, an interaction between Beat Condition and Block was observed (*β* = −0.813, *SE* = 0.115, *z* = −7.081, *p* < 0.001), indicating that the effect of Beat Condition was drastically reduced in the audio-only control block. No other effects or interactions were statistically significant.

#### Experiment 2

All shadowed productions were manually evaluated and annotated for syllable onsets and offsets. We excluded 11 trials (<0.4%) because no speech was produced, and another 232 trials (6.2%) because participants produced more phonemes than the original target pseudoword (e.g., hearing *wasol* but repeating back *kwasol*) which would affect our duration measurements. Thus, acoustic analyses only involved correctly shadowed productions (*n* = 2997; 80%) and incorrectly shadowed productions but with the same number of phonemes (*n* = 504; 13.5%). Measurements of mean F0, duration, and amplitude were calculated for each individual syllable using Praat, and are summarized in Figures S3, S4, and S5 in the Supplementary Material.

Individual syllable measurements of F0 (in Hz), duration (in ms), and amplitude (in dB) were entered into separate Linear Mixed Models as implemented in the lmerTest library (version 3.1-0) in R. Statistical significance was assessed using the Satterthwaite approximation for degrees of freedom ^92^. The structure of the three models was identical: they included fixed effects of Syllable (categorical predictor; with the first syllable mapped onto the intercept), Continuum Step (continuous predictor; scaled and centered around the mean), and Beat Condition (categorical predictor; with a beat gesture on the first syllable mapped onto the intercept), and all interactions. The model also included Participant and Pseudoword as random factors, with by-participant and by-item random slopes for all fixed effects. Model output is given in Table 1 and the combined predicted estimates of the three models are visualized in Figure 2.

**Table 1.**
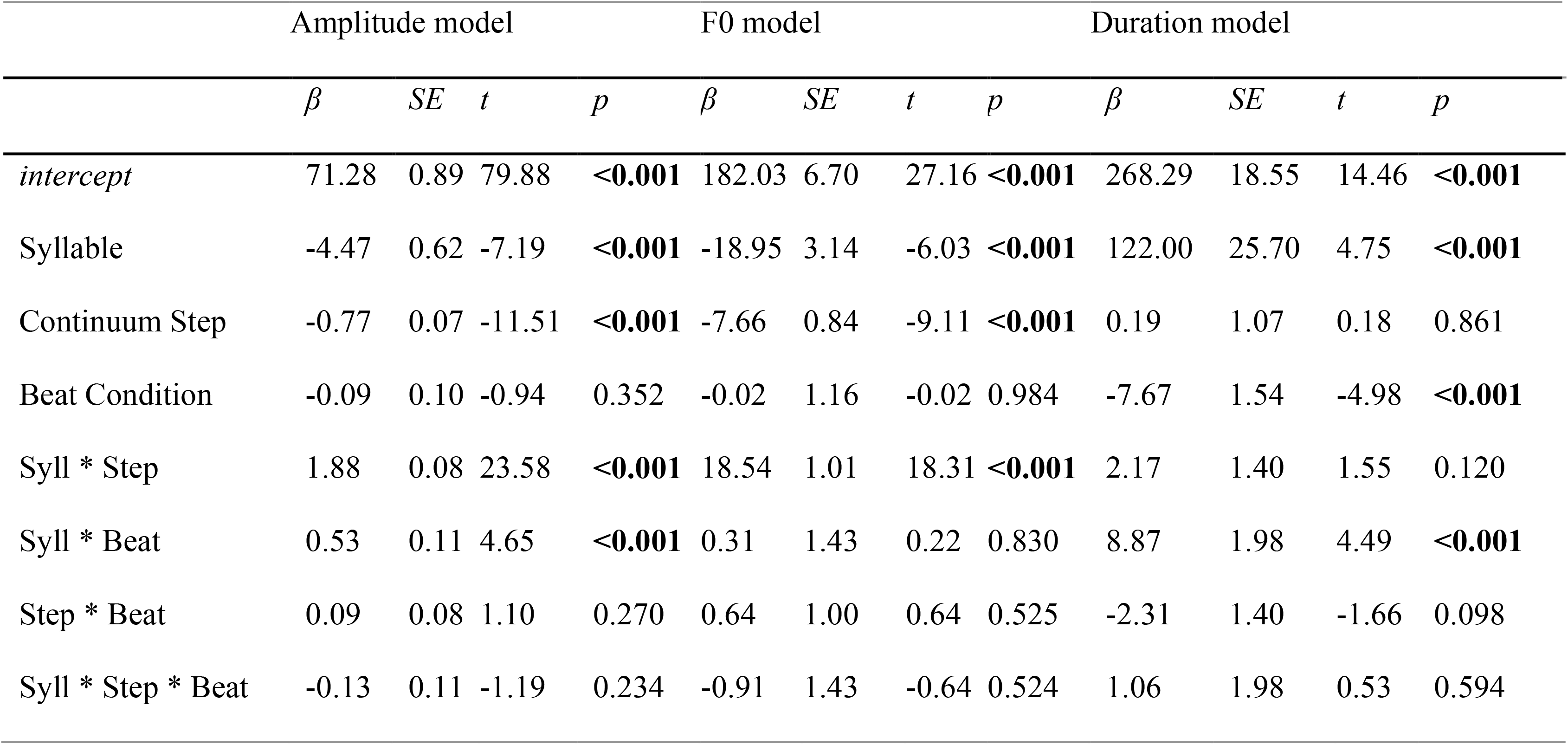
Statistical outcomes of the models testing amplitude (dB), F0 (Hz), and duration (ms) measurements from Experiment 2. Significant effects are highlighted in **bold**.

|  | Amplitude model |  |  |  | F0 model |  |  |  | Duration model |  |  |  |
| --- | --- | --- | --- | --- | --- | --- | --- | --- | --- | --- | --- | --- |
| | $\beta$ | $SE$ | $t$ | $p$ | $\beta$ | $SE$ | $t$ | $p$ | $\beta$ | $SE$ | $t$ | $p$ |
| *intercept* | 71.28 | 0.89 | 79.88 | **<0.001** | 182.03 | 6.70 | 27.16 | **<0.001** | 268.29 | 18.55 | 14.46 | **<0.001** |
| Syllable | -4.47 | 0.62 | -7.19 | **<0.001** | -18.95 | 3.14 | -6.03 | **<0.001** | 122.00 | 25.70 | 4.75 | **<0.001** |
| Continuum Step | -0.77 | 0.07 | -11.51 | **<0.001** | -7.66 | 0.84 | -9.11 | **<0.001** | 0.19 | 1.07 | 0.18 | 0.861 |
| Beat Condition | -0.09 | 0.10 | -0.94 | 0.352 | -0.02 | 1.16 | -0.02 | 0.984 | -7.67 | 1.54 | -4.98 | **<0.001** |
| Syll * Step | 1.88 | 0.08 | 23.58 | **<0.001** | 18.54 | 1.01 | 18.31 | **<0.001** | 2.17 | 1.40 | 1.55 | 0.120 |
| Syll * Beat | 0.53 | 0.11 | 4.65 | **<0.001** | 0.31 | 1.43 | 0.22 | 0.830 | 8.87 | 1.98 | 4.49 | **<0.001** |
| Step * Beat | 0.09 | 0.08 | 1.10 | 0.270 | 0.64 | 1.00 | 0.64 | 0.525 | -2.31 | 1.40 | -1.66 | 0.098 |
| Syll * Step * Beat | -0.13 | 0.11 | -1.19 | 0.234 | -0.91 | 1.43 | -0.64 | 0.524 | 1.06 | 1.98 | 0.53 | 0.594 |

The observed effects of Syllable in all three models demonstrate word-final effects: the last syllables of words typically have lower amplitude, lower F0, and longer durations (cf. Figure 2). The effects of Continuum Step confirm that participants adhered to our instructions to carefully shadow the pseudowords. That is, for higher steps on the F0 continuum (i.e., sounding more WS-like), the amplitude and F0 of the first syllable (mapped onto the intercept) decreased (see simple effects of Continuum Step), while amplitude and F0 of the second syllable increased (interactions Syllable and Continuum Step). Finally, and most critically, the duration of the first syllable (mapped onto the intercept) decreased when the beat gesture was on the second syllable (simple effect of Beat Condition; cf. Figure 2). That is, when the beat fell on the second syllable, the first syllable was more likely to be perceived as unstressed, leading participants to reduce the duration of the first syllable of their shadowed production. Moreover, when the beat fell on the second syllable, the second syllable itself had a higher amplitude and a longer duration (interactions Syllable and Beat Condition; cf. Figure 2). That is, with the beat gesture on the second syllable, the second syllable was more likely to be perceived as stressed, resulting in louder and longer second syllables in the shadowed productions.

#### Experiment 3

Similar to Experiment 1, the categorization responses (SW = 1; WS = 0) were statistically analyzed using a GLMM with a logistic linking function. Fixed effects were Continuum Step and Beat Condition (coding adopted from Experiment 1), and their interaction. The model included random intercepts for Participant, Talker, and Pseudoword, with by-participant, by-talker, and by-item random slopes for all fixed effects.

The model showed a significant effect of Continuum Step (*β* = −0.985, *SE* = 0.162, *z* = −6.061, *p* < 0.001), indicating that higher continuum steps led to lower proportions of SW responses. Critically, it showed a small effect of Beat Condition (*β* = 0.232, *SE* = 0.078, *z* = 2.971, *p* = 0.003), indicating that – in line with our hypothesis – listeners were biased towards perceiving lexical stress on the first syllable if the shadowed production was produced in response to an audiovisual pseudoword with a beat gesture on the first syllable. The interaction between Continuum Step and Beat Condition was marginally significant (*β* = 0.110, *SE* = 0.059, *z* = 1.886, *p* = 0.059), suggesting a tendency for the more ambiguous steps to show a larger effect of the beat gesture.

#### Experiment 4

We analyzed the binomial categorization data (/a:/ coded as 1; /ɑ/ as 0) using a GLMM with a logistic linking function. Fixed effects were Continuum Step and Beat Condition (coding adopted from Experiment 1), and their interaction. The model included random intercepts for Participant and Pseudoword, with by-participant and by-item random slopes for all fixed effects.

The model showed a significant effect of Continuum Step (*β* = −0.922, *SE* = 0.091, *z* = −10.155, *p* < 0.001), indicating that higher continuum steps led to lower proportions of long /a:/ responses. Critically, it showed a small effect of Beat Condition (*β* = −0.240, *SE* = 0.105, *z* = −2.280, *p* = 0.023), indicating that – in line with our hypothesis – listeners were biased towards perceiving the first vowel as /a:/ if there was a beat gesture on the second syllable (or perceiving /ɑ/ if the beat gesture fell on the first syllable). No interaction between Continuum Step and Beat Condition was observed (*p* = 0.817).

## DATA AVAILABILITY STATEMENT

The experimental data of this study will be made publicly available for download upon publication. At that time, they may be retrieved from https://osf.io/b7kue under a CC-By Attribution 4.0 International license.

## ACKNOWLEDGEMENTS

This research was supported by the Max Planck Society for the Advancement of Science, Munich, Germany (H.R.B.; D.P.) and the Netherlands Organisation for Scientific Research (D.P.; Veni grant 275-89-037). We would like to thank Giulio Severijnen for help in creating the pseudowords of Experiments 1-3, and Nora Kennis, Esther de Kerf, and Myriam Weiss for their help in testing participants and annotating the shadowing recordings.

