## Supplementary Information for "Beat gestures influence which speech sounds you hear"

**This Supplementary Information file includes:**

- Supplementary text
  - Experiment S1, replicating Experiment 3
    - including Figure S1
  - Experiment S2, replicating Experiment 4
    - including Figure S2
  - Supplementary figures of Experiment 2,
    - including Figures S3, S4, and S5
- Supplementary table
  - Table S1

Supplementary text

### EXPERIMENT S1

Experiment S1 involved an identical replication of Experiment 3: participants in Experiment S1 were presented with the audio-only shadowed productions, recorded in Experiment 2, and categorized them as having lexical stress on the first syllable (‘strong-weak’; SW) or the last syllable (‘weak-strong’; WS). If beat gestures influence how participants shadowed the audiovisual talker in Experiment 2, then we would predict a higher proportion of SW responses for shadowed productions that followed audiovisual stimuli with a beat gesture on the first syllable.

#### Methods

The method of Experiment S1 was identical to that of Experiment 3. The only difference was that a new sample of twenty-six participants was recruited (22 females, 4 males; mean age = 24, range = 18-34) from the Max Planck Institute for Psycholinguistics participant pool.

#### Results

The categorization data, calculated as the proportion of SW responses, are presented in Figure S1. The overall patterns are very similar to Experiment 3. Colors indicate whether the original audiovisual pseudoword the talkers were instructed to shadow was produced with a beat gesture on the first (blue/darkgray) or the second syllable (orange/lightgray).

Like in Experiment 3, the blue (darkgray) line seems to show a higher proportion of SW responses than the orange (lightgray) line, though the effect is small.

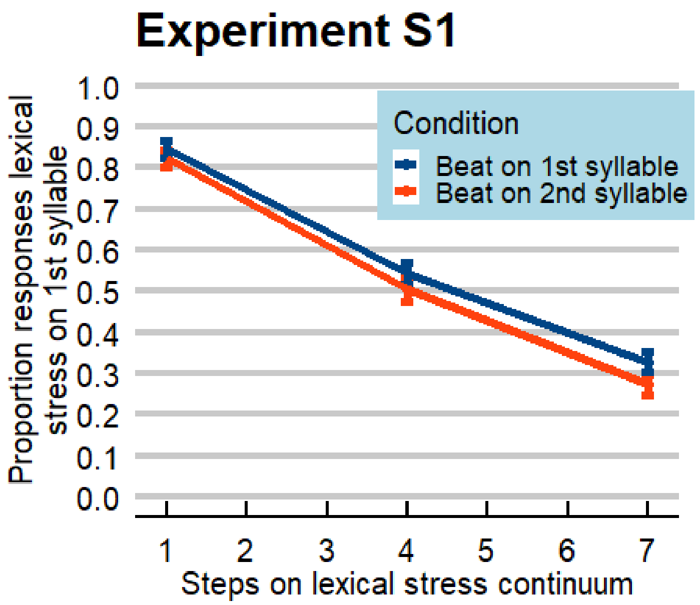


**Figure S1. Results from Experiment S1, replicating Experiment 3.** Proportion of trials in which participants reported perceiving lexical stress on the first syllable (e.g., *WAsol* vs. *waSOL*). Participants heard audio-only shadowed productions from Experiment 2 which themselves were produced after audiovisual stimuli, sampled from a lexical stress continuum varying F0 independently for the two syllables. If the audiovisual stimulus in Experiment 2 involved a pseudoword with a beat gesture on the first syllable, the shadowed production was more likely to be perceived as having lexical stress on the first syllable (SW; blue/darkgray). Conversely, if the audiovisual stimulus in Experiment 2 involved a pseudoword with a beat gesture on the second syllable, the shadowed production was more likely to be perceived as having lexical stress on the second syllable (WS; orange/lightgray). Error bars enclose 1.96 x SE on either side; that is, the 95% confidence intervals.

A GLMM with identical structure to the one used in Experiment 3 showed a significant effect of Continuum Step ($\beta$ = -1.276, *SE* = 0.171, $z$ = -7.468, $p$ < 0.001), indicating that higher continuum steps led to lower proportions of SW responses. Critically, it showed a small effect of Beat Condition ($\beta$ = 0.213, *SE* = 0.083, $z$ = 2.562, $p$ = 0.010), replicating Experiment 3. That is, listeners were biased towards perceiving lexical stress on the first syllable if the shadowed production was produced in response to an audiovisual pseudoword with a beat gesture on the first syllable. The interaction between Continuum Step and Beat Condition was, like in Experiment 3, marginally significant ($\beta$ = 0.102, *SE* = 0.062, $z$ = 1.655, $p$ = 0.098), suggesting a tendency for the more ambiguous steps to show a larger effect of the beat gesture.

### EXPERIMENT S2

Experiment S1 involved an identical replication of Experiment 4: participants in Experiment S2 were presented with audiovisual stimuli in which a talker produced a beat gesture on either the first or the last syllable of disyllabic sentence-final target pseudowords. Pseudowords were manipulated to be ambiguous in lexical stress ('average' values of F0, amplitude, and duration) and the vowel of the first syllable was sampled from F2 continua, varying from long /a:/ to short /ɑ/.

#### Methods

The method of Experiment S2 was identical to that of Experiment 4, except that new participants were tested. Specifically, the same participants recruited for Experiment S1 also performed Experiment S2 in a fixed order: first Experiment S2, then S1.

**Results**

The proportions of long /a:/ responses are presented in Figure S2, which looks very much like the data from Experiment 4. Once again, there seems to be a small difference between the blue/darkgray (beat gesture aligned to first vowel onset) and orange/lightgray line (beat gesture aligned to second vowel onset), indicative of an influence of the visually presented beat gesture. Critically, the blue/darkgray line seems to show a lower proportion of long /a:/ responses than the orange/lightgray line.

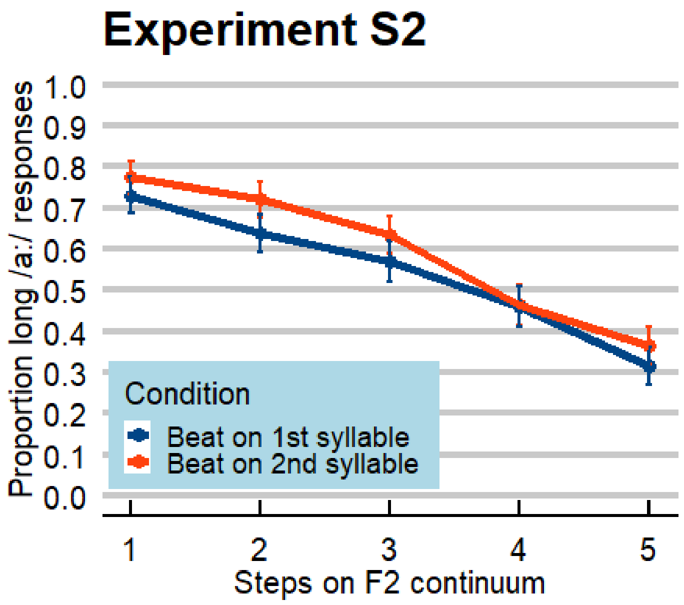


**Figure S2. Results from Experiment S2, replicating Experiment 4.** Proportion of trials in which participants reported perceiving a long /a:/ as first vowel in the sentence-final pseudowords (e.g., *baagpif* vs. *bagpif*). Each target word contained a first vowel that was ambiguous between /ɑ/ and /a:/ (fixed vowel duration and F1, varying F2 on a 5-step continuum). Moreover, prosodic cues to lexical stress (F0, amplitude, syllable duration) were set to ambiguous values. Participants were more likely to report perceiving a long /a:/ as the first target vowel if the speaker produced a beat gesture on the second syllable (in orange/lightgray), presumably because the ambiguous vowel duration was relatively long for an unstressed syllable. Conversely, the proportion of long /a:/ responses was lower if the speaker produced a beat gesture on the first syllable (in blue/darkgray), because the ambiguous vowel duration was relatively short for a stressed syllable. Error bars enclose 1.96 x SE on either side; that is, the 95% confidence intervals.

A GLMM with identical structure as used in Experiment 4 showed a significant effect of Continuum Step ($\beta$ = -0.867, *SE* = 0.116, $z$ = -7.475, $p$ < 0.001), indicating that higher continuum steps led to lower proportions of long /a:/ responses. Critically, it showed a small effect of Beat Condition ($\beta$ = -0.257, *SE* = 0.125, $z$ = -2.050, $p$ = 0.040), indicating that – in line with our hypothesis – listeners were biased towards perceiving the first vowel as /a:/ if there was a beat gesture on the second syllable (or perceiving /ɑ/ if the beat gesture fell on the first syllable). No interaction between Continuum Step and Beat Condition was observed ($p$ = 0.622).

**SUPPLEMENTARY FIGURES OF EXPERIMENT 2**

The figures below summarize the average amplitude (Figure S3), fundamental frequency (F0; Figure S4), and duration values (Figure S5) of the shadowed productions from Experiment 2.

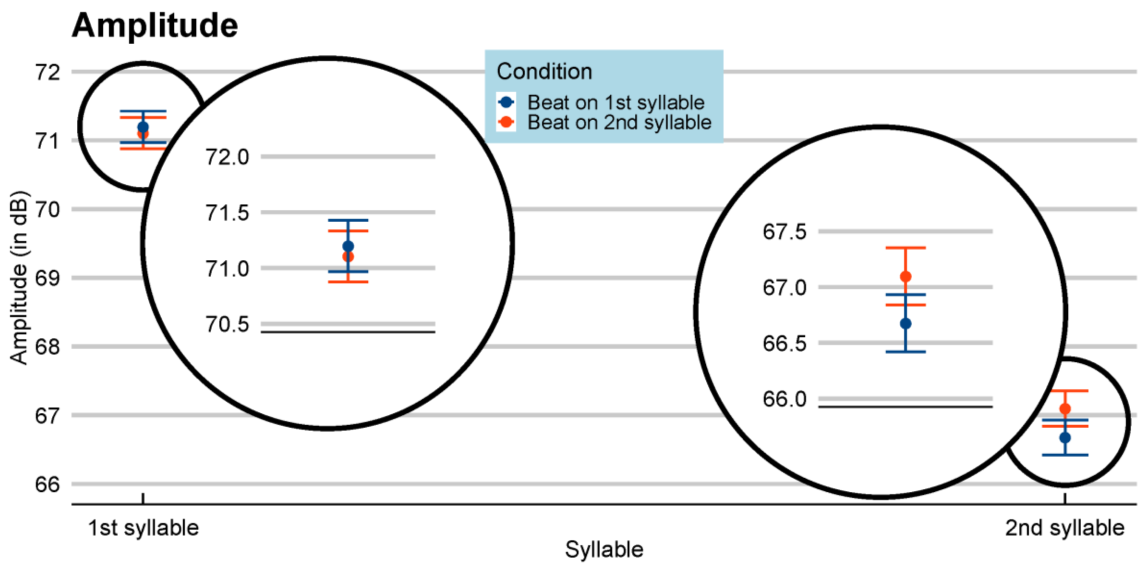


**Figure S3. Average amplitude values (dB) of the shadowed productions from Experiment 2.** Values are averaged over continuum steps, plotted separately for the first syllable (left) and the second syllable (right), split for shadowed productions following an audiovisual trial with a beat gesture on the first syllable (in blue/darkgray) vs. on the second syllable (orange/lightgray). The overall data show a pronounced word-final effect, with lower amplitude for second syllables vs. first syllables. The insets demonstrate a small but significant effect of Beat Condition in the second syllable: if the audiovisual talker produced a beat gesture on the second syllable, shadowers produced the second syllable with approximately 0.5 dB higher amplitude. Error bars enclose 1.96 x SE on either side; that is, the 95% confidence intervals.

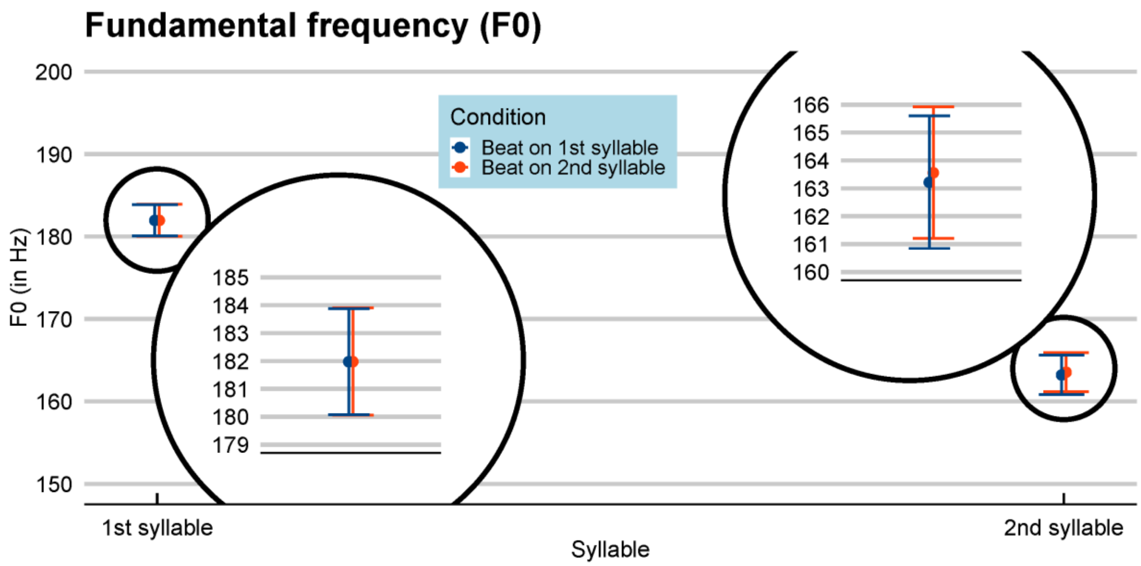


**Figure S4. Average fundamental frequency values (F0; in Hz) of the shadowed productions from Experiment 2.** Values are averaged over continuum steps, plotted separately for the first syllable (left) and the second syllable (right), split for shadowed productions following an audiovisual trial with a beat gesture on the first syllable (in blue/darkgray) vs. on the second syllable (orange/lightgray). The overall data show a pronounced word-final effect, with lower F0 for second syllables vs. first syllables. The insets demonstrate the fact that no significant effect of Beat Condition was found in the F0 data. Error bars enclose 1.96 x SE on either side; that is, the 95% confidence intervals.

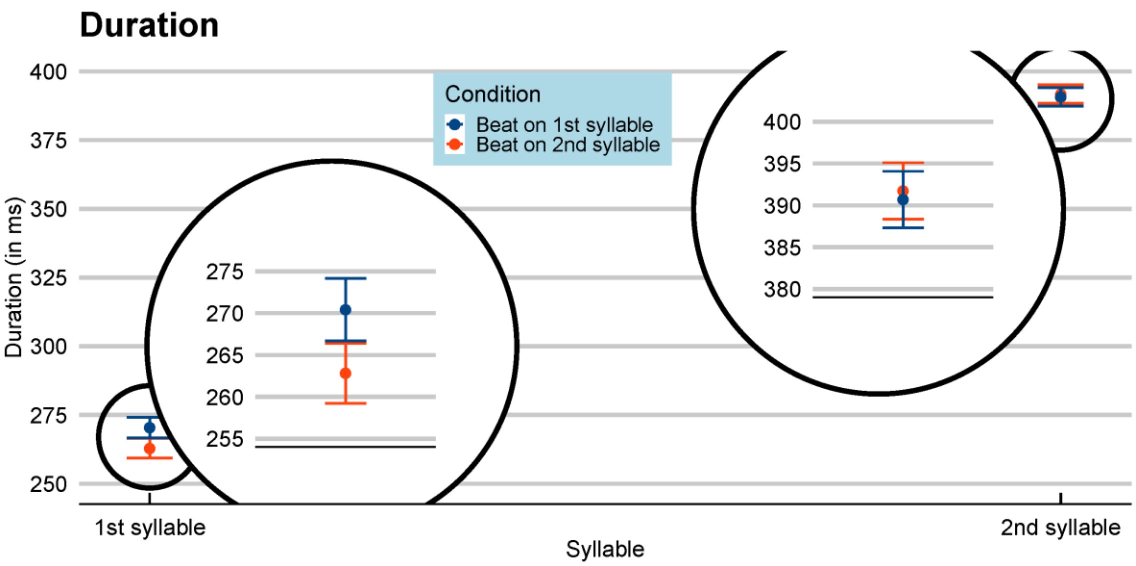


**Figure S5. Average duration values (ms) of the shadowed productions from Experiment 2.** Values are averaged over continuum steps, plotted separately for the first syllable (left) and the second syllable (right), split for shadowed productions following an audiovisual trial with a beat gesture on the first syllable (in blue/darkgray) vs. on the second syllable (orange/lightgray). The overall data show a pronounced word-final effect, with longer durations for second syllables vs. first syllables. The insets demonstrate a small but significant positive effect of Beat Condition in the first syllable, and a small but significant negative effect of Beat Condition in the second syllable. That is, if the audiovisual talker produced a beat gesture on the first syllable, shadowers increased the duration of their first syllable by about 8 ms (compared to seeing a beat gesture on the second syllable). Error bars enclose 1.96 x SE on either side; that is, the 95% confidence intervals.

### Supplementary table

**Table S1. List of pseudowords (in IPA) used in the experiments.** The question mark in the pseudowords of Experiment 4 indicates the location of the 5-step vowel continua varying F2 from short /ɑ/ to long /a:/.

| | **pseudowords Experiments 1-3** | **pseudowords Experiment 4** |
|---|---|---|
| 1 | /ʋa.sɔl/ | /bʔx.pɪf/ |
| 2 | /ɛl.pɑt/ | /pʔf.bʏx/ |
| 3 | /kla.fɔs/ | /bʔx.kʏf/ |
| 4 | /lo.sɛp/ | /tʔx.tɔs/ |
| 5 | /nu.fa/ | /tʔf.pɛx/ |
| 6 | /plo.sɪm/ | /tʔf.dɔs/ |
| 7 | /pra.bɔp/ | /pʔg.dɔx/ |
| 8 | /rɔs.kɪl/ | /bʔf.kix/ |
| 9 | /rʏŋ.ka/ | |
| 10 | /stra.dɔt/ | |
| 11 | /tu.sa/ | |
| 12 | /dɛm.rɔf/ | |
